# WeavePop: A bioinformatics workflow to explore and analyze genomic variants of eukaryotic populations

**DOI:** 10.1101/2025.08.15.670593

**Authors:** Claudia Zirión-Martínez, Paul M. Magwene

## Abstract

Analyzing genomic variants in large datasets composed of short-read sequencing data is a process that requires multiple steps and computational tools, which makes it a complicated task that is difficult to reproduce across projects and laboratories. To address this need, we developed a reproducible and scalable Snakemake workflow called WeavePop, which aligns samples to selected references, obtains reference-based assemblies, annotations, and sequences, and identifies small variants and copy-number variants in eukaryotic haploid organisms. All the results are integrated into a database that can be easily shared and explored through a graphical web interface provided alongside the workflow, making the discovery of variants in a population of study very simple. WeavePop is available from GitHub (https://github.com/magwenelab/WeavePop) for Linux operating systems. Here we exemplify the use of WeavePop in a large collection of isolates of the pathogenic fungus *Cryptococcus neoformans*.

**Graphical abstract:** 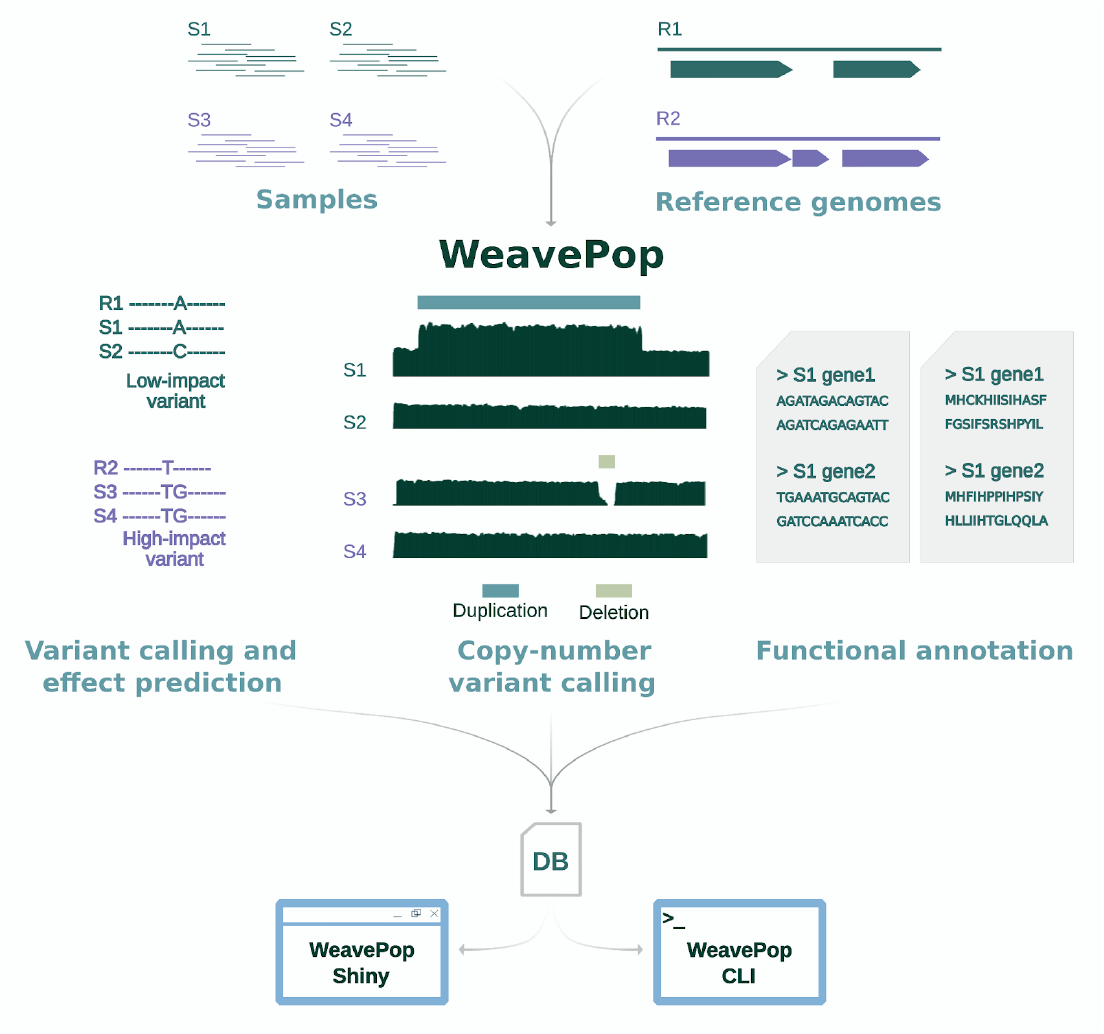

## Introduction

Sequencing whole genomes, particularly those of microbial organisms, has become a routine and relatively inexpensive procedure for many investigators and research teams. It is not unusual for a single research group to sequence tens to hundreds of genomes annually for various purposes, including population genomics, experimental evolution, mutational screens, and phylogenomics. In addition, for many model organisms, hundreds or thousands of whole-genome sequencing runs are available in public databases such as the National Library of Medicine Sequence Read Archive and the European Nucleotide Archive. Despite the ubiquity of genome sequence data, the computational process required to go from raw sequence reads to mapped genomes while also detecting both small-scale and structural variation remains a significant challenge due to the number of steps it involves and the complexity of the tools required. Moreover, ensuring that this process is reproducible and the resulting data is accessible to users with varying levels of computational expertise adds greater complications to the task.

To address this computational hurdle, we developed WeavePop (**W**orkflow to **E**xplore and **A**nalyze **V**ariants of **E**ukaryotic **Pop**ulations), a bioinformatics pipeline that carries out an extensive set of analyses to provide complementary information about genomic variation within a collection of eukaryotic haploid organisms. WeavePop detects and catalogs large-scale and small-scale variation by including reference-based read mapping (allowing for multiple references), variant calling, variant effect prediction, identification of repetitive sequences, and copy-number variant (CNV) detection. To accomplish these tasks, WeavePop takes advantage of multiple field-standard tools in combination with new algorithms, bringing them together in a pipeline that facilitates their application to multiple use cases. This saves time and generates a diverse set of outputs that are convenient to evaluate directly or that can be passed to downstream analyses.

### anaconda

WeavePop is implemented as a Snakemake (Mölder et al. 2021) workflow. The use of Snakemake to implement this pipeline makes it readable, scalable, reproducible, and easy to configure and execute. WeavePop is available through GitHub for Linux operating systems, and all of the required software tools are readily installed using predefined Conda environments (https://anaconda.com) managed by Snakemake.

The numerous outputs generated by WeavePop across multiple samples are combined into a single DuckDB (Raasveldt and Mühleisen 2019) SQL database file, which can be readily shared, archived, and used to explore the output in a user-friendly manner. This database can be accessed via standard SQL syntax or using DuckDB client APIs for popular languages such as Python and R. In addition, we provide a local Shiny (https://shiny.posit.co/py/) web application that facilitates querying the database for variants and their predicted effects, CNVs, functional annotation, and gene sequences. In sum, using the databases produced by WeavePop allows a single lab or small research community to produce easily shared resources that facilitate exploration and analysis of genomic variation.

Here we discuss the structure and implementation of WeavePop. We then illustrate the utility of WeavePop by applying it to study genome variation in a large collection (1,025 samples) of *Cryptococcus neoformans. C. neoformans* is a fungal pathogen of worldwide concern for human health (Brown et al. 2024), and we illustrate the application of WeavePop to detect and catalog population genomic structural variation in this species.

## Methods

### Overview

WeavePop aims to detect and catalog genomic variation between a set of samples and one or more suitable reference genomes. To exploit the availability of multiple reference genomes, WeavePop allows each sample to be aligned to a different reference genome, such as lineage-specific whole genome assemblies, while at the same time facilitating comparisons across samples regardless of reference. For each sample, the workflow produces a read alignment, a reference-based assembly along with its functional annotation, the sequence of all genes, small variants (single and multiple nucleotide polymorphisms and indels) along with their predicted effects, and CNVs. An overview of the workflow is shown in Figure 1, and the Directed Acyclic Graph (DAG) of Snakemake tasks used to implement this workflow is shown in Figure S1.

**Figure 1.**
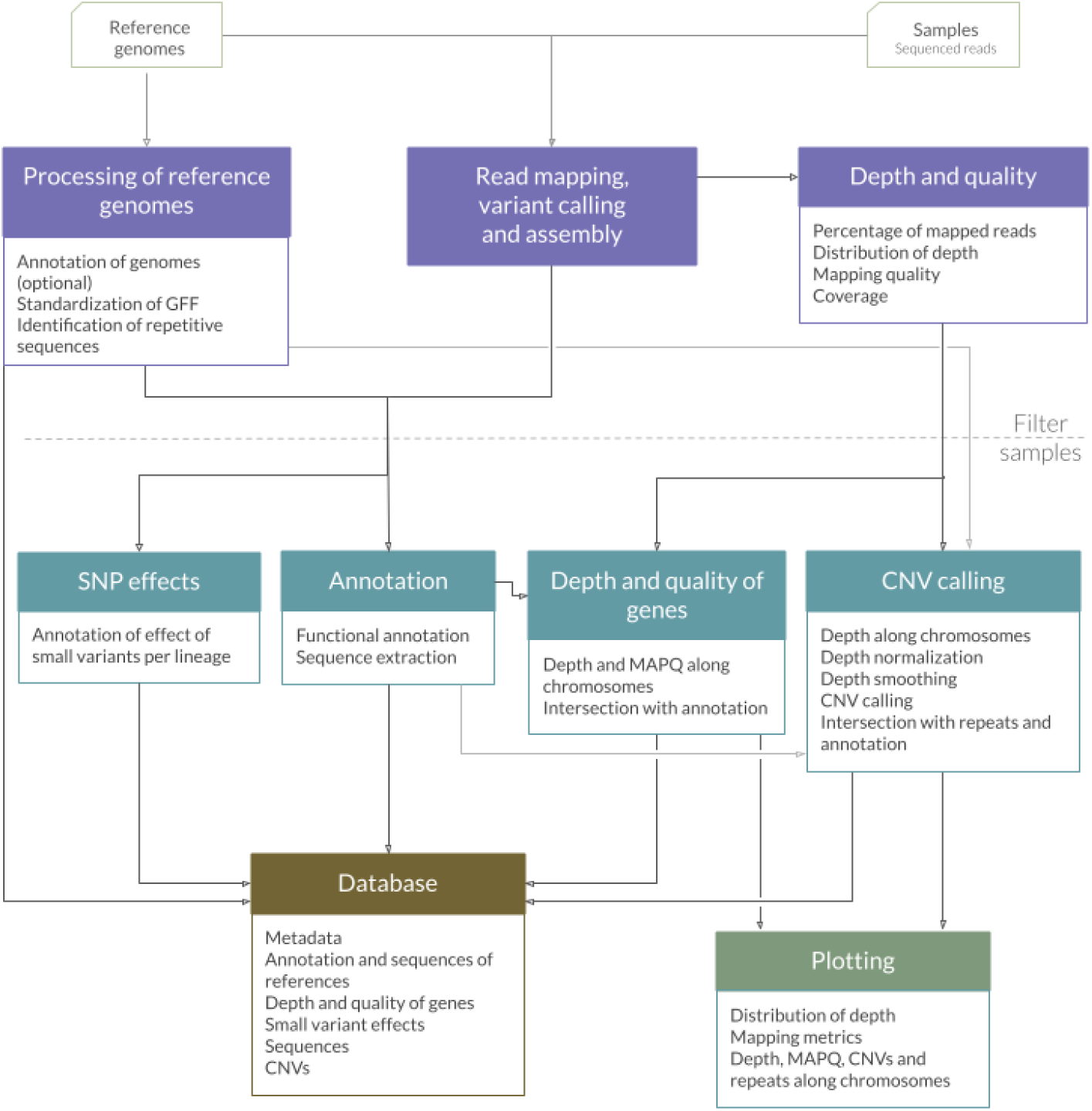
Overview of the modules in WeavePop. The required modules, shown in purple, are for processing the reference genomes, read mapping, variant calling, and the analysis of the depth and quality. An optional step of filtering out low-quality samples can be included before the next steps. The optional modules, shown in blue, are for the annotation of SNP effects, functional annotation, the intersection of mapping quality (MAPQ) and depth with the annotated features, and copy-number variant (CNV) calling. The database module integrates the results of the previous modules into an SQL database. The plotting module generates graphs for each sample and summary graphs of the final dataset. The activation of a module automatically activates the required steps in the previous ones.

### Snakemake description

WeavePop is implemented in Snakemake, a workflow management system written in Python, that organizes the execution of the steps required to generate target outputs. Snakemake facilitates both parallelization and efficiency. If the user wishes to re-execute the workflow with changes in the configuration and parameters, or if the workflow stops after a failure, Snakemake only repeats the execution of the steps whose results would be updated, saving time and storage. Snakemake also makes it possible to generate reproducible analyses because all the configurations are documented, and all the software tools used in multi-step analysis are contained in Conda environments managed by Snakemake. After downloading WeavePop and installing Conda, the user only needs to run the following command to install Snakemake.

~~~
conda env create --file workflow/envs/snakemake.yaml
~~~

### Input

The pipeline requires each sample’s paired-end short-read FASTQ files and a reference genome or genomes (see below the description of Module 1). The relationship between each sample and a reference genome is provided in a metadata table (File S1).

A table that matches the chromosome sequence IDs of each reference genome with its conventional name is needed. This allows the comparison of chromosomes across reference genomes (File S2).

The repetitive sequences of the reference genomes are identified using RepeatModeler (Flynn et al. 2020) and RepeatMasker (https://www.repeatmasker.org/), which requires a database (FASTA file) of known repetitive sequences, such as the RepBase (Bao et al. 2015). The user can provide this file or allow WeavePop to use a mock database if an accurate identification is not needed.

Finally, two optional files can be provided: one to highlight the location of genes or genetic features of interest in the plots (File S3) and a list of sample names to exclude from all analyses.

### Configuration

All the configuration options required to complete a run of WeavePop are specified in a single file (File S4). The paths to the input files and output directory, as well as the parameters for the tools, are set in this file. The modular structure of the workflow permits the process to be run step by step. If no modules are activated, only the mandatory steps will be executed (Module 1, except for the identification of repetitive sequences, and Modules 2 and 3). With these, one can evaluate the quality of the alignments and decide to adjust the filtering parameters (see Module 3 below) before continuing. Modules 4 to 7 can be activated independently. To run every module of the analysis, only the database and the plotting modules need to be explicitly activated. Besides the modules mentioned above (analysis workflow), one can choose to integrate the results of multiple datasets into one (see the Combining datasets section below).

The command-line options for Snakemake can be set in a YAML file that specifies the execution profile. The default profile provided contains the settings needed by the workflow (File S5).

After editing the configuration file and execution profile, the user only needs to run the following to execute WeavePop:

~~~
conda activate snakemake
snakemake --profile config/default
~~~

### Modules

The nine modules of the WeavePop workflow, including the tasks they accomplish and the tools employed, are described below.

#### 1. Processing of reference genomes

This module provides scripts to standardize the format of the reference genome annotation GFF files, homogenize the nomenclature of feature IDs, and add intron and intergenic features to the annotation. These steps are implemented using AGAT (Dainat 2022) and Python. RepeatModeler and RepeatMasker are used to identify repetitive sequences in the reference genomes.

For this module, the inputs are a FASTA file and a GFF annotation file for each reference genome. Alternatively, if homogeneous annotation for all references is desired or there are no annotations available for all references, the user can provide a single GFF file, which will be used to annotate the rest of the references via Liftoff (Shumate and Salzberg 2021). Doing so facilitates the comparison of variation in genic regions and other annotated features even when samples are aligned to different references.

#### 2. Read-mapping, variant calling, and assembly

Snippy (Seemann 2020), a haploid variant calling pipeline, performs the central analysis of the WeavePop workflow, using the paired-end short reads of each sample and the corresponding reference genome. Snippy uses BWA (Li and Durbin 2009) to align the reads to the reference and FreeBayes (Garrison and Marth 2012) to call small variants. It also generates a reference-based assembly, which is a version of the reference genome with the called variants instantiated.

#### 3. Depth and quality

To analyze the quality of the read alignments of each sample, several metrics of read depth, mapping quality (MAPQ), and coverage are obtained in this module. Samtools (Li et al. 2009) is used to filter the BAM file with a user-defined threshold (`min_mapq`) for the MAPQ of the aligned reads. The `stats` command from Samtools is then used to get the read depth distribution of each chromosome from the raw and filtered BAM files. From the distributions, the mean and median depth of each chromosome and the genome-wide mean and median are obtained. Samtools `stats` is also used to obtain, from the raw BAM file, the percentage of unmapped, mapped, and properly paired reads, along with the percentage of reads with low, intermediate, and high MAPQ, delimited by user-defined thresholds. The `coverage` command of Samtools is used to get the coverage of the raw and filtered reads. These metrics are used to flag samples according to user-defined thresholds: minimum percentages of high-quality alignments and mapped reads, coverage, and genome-wide median depth. The flagged samples can be excluded from all further steps.

Finally, to estimate the mean read depth of windows of a fixed size (specified by the user) along each chromosome, to be used in modules 6 and 7, Mosdepth (Pedersen and Quinlan 2018) is used on the high-quality reads.

#### 4. Annotation of SNP effects

This module generates predicted coding and modifier effects of small variants. The VCF files of the samples that were aligned to a reference are compiled, using the Bcftools `isec` command (Danecek et al. 2021), to obtain a VCF of all the variants in that group of samples, along with a presence/absence matrix of the variants in each sample. The new VCF file is then annotated with SnpEff (Cingolani et al. 2012) using the functional annotation of the corresponding reference. The annotations in the VCF are then transferred to multiple tables.

#### 5. Annotation

Liftoff annotates each sample’s reference-based assembly using the annotation of the corresponding reference. Then AGAT adds the intergenic and intronic features to the GFF files and extracts the sequences of the annotated transcripts of all genes. This produces a FASTA file per sample with all the predicted nucleotide coding sequences and another with the corresponding amino acid sequences.

#### 6. Depth and quality of genes

This module implements windowed analyses of mapped read depth and quality using the Mosdepth results from Module 3, and Samtools `mpileup` along with Bedtools (Quinlan and Hall 2010) to estimate the mean MAPQ of the windows. Per-window metrics of read depth and quality are intersected with the annotation to estimate the mean depth and mean MAPQ of each annotated genomic feature, including each gene and its nested features.

#### 7. Copy-number variant calling

In this module, CNVs are identified using the read depth estimates obtained with Mosdepth in Module 3. The median depth of all the windows in the genome is used to normalize the mean depth of each window along the chromosomes. The normalized read-depth is then smoothed using SciPy’s ndimage median filter algorithm (Virtanen et al. 2020) with a smoothing size specified by the user. Bedtools is used to calculate the fraction of each window that overlaps with repetitive sequences of the corresponding reference genome (identified in Module 1). Each window is classified as deleted, duplicated, or single-copy using a user-defined threshold based on the smoothed normalized read-depth values. Consecutive windows with the same classification are fused into regions, and duplicated/deleted regions are catalogued in a table. A second table summarizing the full set of CNVs for each chromosome is also generated, with metrics such as the number of regions in the chromosome and the percentage of chromosomal length covered by each CNV type.

#### 8. Database

The main outputs of the workflow are combined into a relational database to facilitate querying and exploration. WeavePop employs the DuckDB database system, a portable and efficient platform that supports a rich SQL dialect and provides APIs for many commonly used programming languages such as R and Python. The DuckDB database that WeavePop creates is a single-file database containing the key tables generated by modules 4, 6, and 7, the sequences of the annotated features, the annotation tables of the reference genomes, plus the user-provided metadata table. The schema of the database can be found in Figure S2.

The WeavePop database can be used to extract results of interest in a simpler and more powerful manner than from the original tables and files alone. Since this is a relational SQL database, multiple pieces of information from different tables can be combined using SQL queries to obtain specific outputs of interest, such as all the genes in a certain chromosomal region that have variants with high-impact effects. Using the database is further simplified using the provided tool, WeavePop-Shiny, a Shiny for Python application. WeavePop-Shiny provides a graphical web interface that helps users unfamiliar with SQL to filter and extract results from the database using predefined functions that generate output tables (Figure 2). For integration with other command-line tools, we also provide a command-line interface, WeavePop-CLI, that implements the same functions as WeavePop-Shiny, while also allowing for arbitrary SQL queries (for the users who know SQL). Examples of the use of WeavePop-CLI are given in File S6.

**Figure 2.**
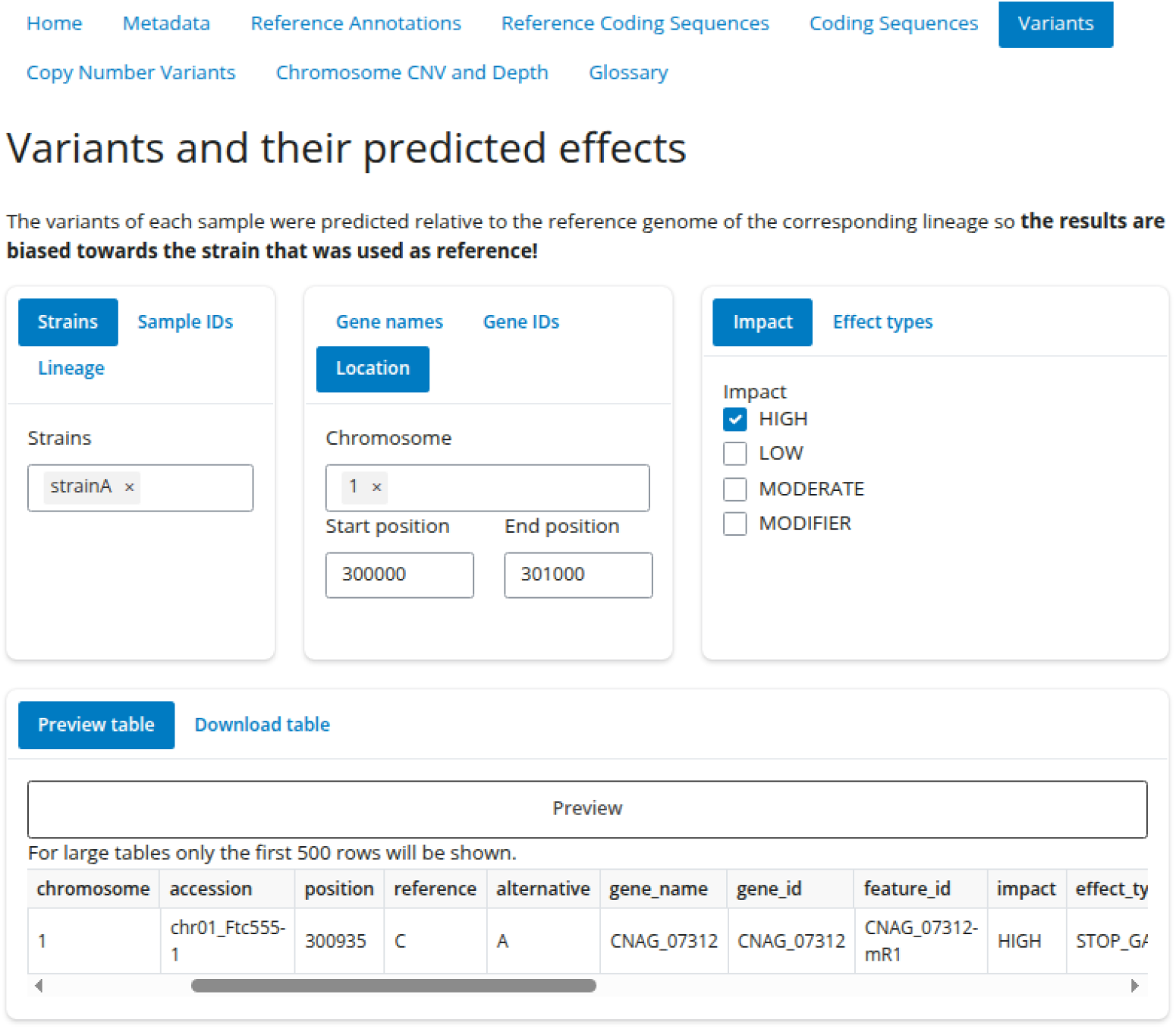
Screenshot of the Variants tab of WeavePop-Shiny. It shows an example of a combination of filters to obtain the high-impact variants in a specific range of coordinates of one strain.

#### 9. Plotting

Results from the modules 1, 3, 6, and 7 are used to create plots that show quality metrics, the read-depth distribution, and the read-depth and MAPQ of each sample along the chromosomes, with the repetitive sequences of the reference and the identified CNV regions. An optional list of gene IDs provided by the user can be used to obtain the coordinates of the features from the reference genome to include user-defined genetic features in these plots (e.g., centromere flanking genes, mating type locus, individual genes of interest, etc.).

### Output

In addition to the DuckDB database, all intermediate and final outputs produced by WeavePop are organized into a single hierarchical directory structure. This directory structure has been structured to allow for flexibility in downstream analyses as well as exploration of alternate tool chains. The directory structure was also designed to minimize additional computation required if a subset of samples is changed or a set of analyses is re-run. A full description of the output files is given in File S7.

### Combining datasets

Recognizing that population genomic resources grow over time, we have implemented facilities for combining the outputs of multiple WeavePop runs. This allows for simple integration of samples from different sequencing projects or data from multiple labs, avoiding redundant computation and storage as more data is obtained.

## Results and Discussion

To demonstrate the use of WeavePop, we re-analyzed the genomes of 1,025 isolates of the basidiomycete fungus *C. neoformans* (Desjardins et al. 2017; Ashton et al. 2019). *C. neoformans* is an opportunistic pathogen and is considered a critical priority pathogen by the World Health Organization (WHO 2022). Here, we demonstrate an analysis of chromosomal structural variation, facilitated by the CNV calling pipeline of WeavePop. Aneuploidy and large-scale copy number variation have been shown to contribute to traits such as resistance to antifungal drugs in *Cryptococcus* and other fungal pathogens (Morrow and Fraser 2013). The brief analysis here focuses on differences in aneuploidy and CNV between the major lineages of *C. neoformans* and between environmental and clinical strains. A more detailed analysis of *C. neoformans* structural variation will be presented elsewhere.

### *Cryptococcus neoformans* aneuploidy and large-scale copy number variation

To identify aneuploidy and large-scale copy number variation in *C. neoformans*, we combined and reanalyzed genome data from two earlier studies on the global genomic diversity of this species. The study by Desjardins et al. (2017) reported genome data for 387 *C. neoformans* strains, representing each of the four major lineages (VNI, VNII, VNBI, VNBII), and including both environmental and clinical isolates. A later study by Ashton et al. (2019) described a sample of 678 isolates of the VNI lineage collected from various countries in Asia. We used the dataset joining facilities of WeavePop to conduct a joint analysis of the strains from these two data sources. After removing 40 samples due to low quality or uncertain baseline ploidy, we analyzed a final dataset of 1,025 isolates (Supplementary Material and Tables S1 and S2, and S3).

We used the results of the WeavePop CNV-calling module to distinguish aneuploid and partially duplicated chromosomes (Supplementary Material and Table S4) in this dataset. None of the 1,025 isolates contains deleted chromosomes, but 36 strains (3.5%), all of clinical origin, have at least one fully duplicated chromosome (Figure 3a, Figure S3). When considering partially duplicated chromosomes (20-80% of their length present in two or more copies), we identified an additional 25 strains, all clinical isolates as well (Figure 3a). In sum, nearly 6% of *C. neoformans* strains have large-scale chromosomal duplications. The chromosomes that we found to be most frequently fully duplicated were chromosomes 12 and 13, which were aneuploid in 12 strains each (Figure 3b and Figure S4). Our finding is consistent with several other studies that identified chromosomes 12 and 13 among the most frequently duplicated chromosomes (Stott et al. 2024; Rhodes et al. 2017). Duplicated copies of chromosomes 1, 4, and 6 were only found in isolates of lineage VNI; chromosome 9 and 12 duplication were only found in samples of lineages VNI and VNBII, and chromosome 13 and 14 in samples of lineages VNI, VNBI, and VNII (Figure 3b). Within the lineage VNI, for which we have the most samples (n = 827), 29 strains had fully duplicated chromosomes (3.5%). Chromosomes 4 and 14 were duplicated in one isolate each; chromosomes 1 and 6 in two, chromosome 9 in 5, chromosome 13 in 8, and chromosome 12 in 11 (Figure 3b).

**Figure 3.**
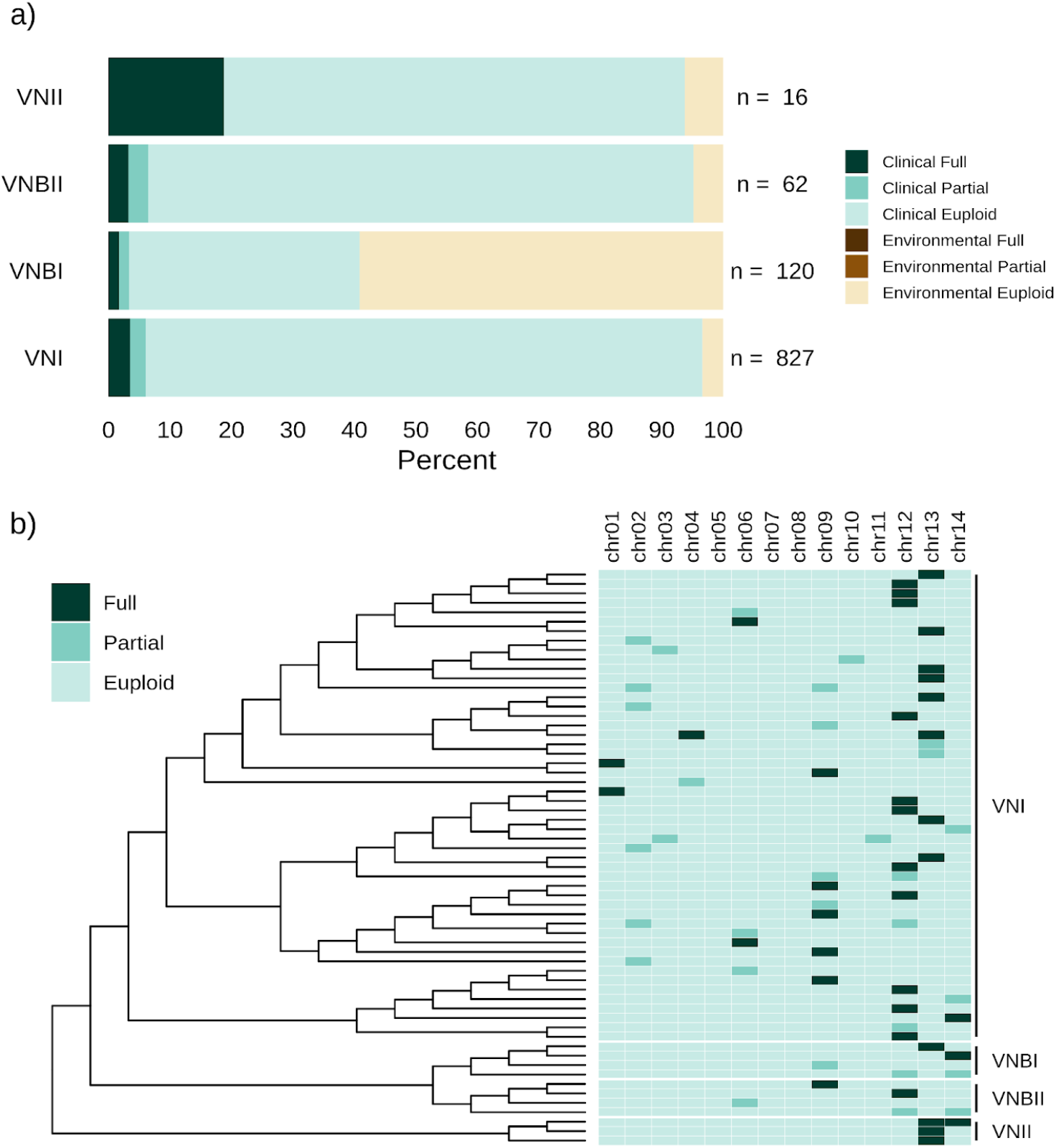
Aneuploidy in *C. neoformans*. a) Percentage of samples per lineage and source of isolation with at least one fully duplicated chromosome (Full: >=80% of chromosome length covered by called duplications; in VNI n=29, in VNII n=3, and VNBI and VNBII n=2), at least one partially duplicated chromosome (Partial: 20-80%; in VNI n=21, in VNBI and VNBII n=2), or no duplication (Euploid: 0-20%). The bars represent the percentage of samples in each category for each lineage, and the numbers to the right of the bars indicate the total number of samples analyzed per lineage. b) Reduced phylogeny of the samples with chromosomes fully or partially duplicated. The right section has one column per chromosome with cells colored by category of duplication.

An important caveat for all large-scale analyses of copy number variation in microbial organisms is the possibility that duplicated chromosomes may be lost with passaging prior to genome sequencing. For example, in *C. neoformans* chromosome 1, aneuploidy has been shown to be selected for under exposure to the antifungal drug fluconazole (Sionov et al. 2010). However, chromosome 1 aneuploidy tends to be highly deleterious in the absence of azole drugs and is typically lost after a modest number of passages on non-selective media. The number of passages that the isolates analyzed here underwent prior to genome sequencing is unknown, and hence we can not rule out the possibility that post-collection selection against aneuploidy biases the chromosomal patterns we detected.

### Related tools

There is an abundance of computational programs that identify genomic variants from short-read sequences and reference genomes. Some are focused on small variants (single and multiple nucleotide polymorphisms and indels), and others are focused on large variants (CNVs and/or complex structural variants). However, to our knowledge, only a few workflows include both kinds of variants.

Similar to WeavePop, PerSVade (Schikora-Tamarit and Gabaldón 2022) identifies both small and large-scale genomic variation, including variant calling with effect prediction, estimated read depth of genes, and large copy-number variants. PerSVade identifies complex structural variants with personalized parameters, making it a more suitable tool for detailed analysis of individual samples. A key difference that gives WeavePop a special purpose is the fact that it can efficiently analyze large numbers of samples and give population-scale results. A second workflow that also detects small and large genomic variation is nf-core/sarek (Hanssen et al. 2024). This tool includes many software options for each step and is designed to work on multiple samples, but it is designed primarily for biomedical research on humans and mice.

In comparison to the available tools, WeavePop has a unique advantage in that it generates a user-friendly and shareable database that facilitates a quick exploration, without the need to manipulate tables or extract information from files of specific formats.

## Conclusion

WeavePop is a reproducible, readable, customizable, and scalable workflow for genomic analysis of haploid eukaryotic populations. It uses field-standard programs to detect small variants and annotate their effect, identify CNVs, and provide reference-based assemblies, functional annotation, and sequences of multiple samples. It generates a database to explore variation in a population, which can be used as a shared resource for research communities. A future version of WeavePop is intended to include the discovery of complex structural variants and the possibility of analyzing diploid genomes.

## Data availability

WeavePop is available at GitHub https://github.com/magwenelab/WeavePop, which includes instructions, examples, and recommendations for users in the repository’s Wiki https://github.com/magwenelab/WeavePop/wiki.

The repository describing the analysis of the Cryptococcus neoformans strain collection using WeavePop is available at https://github.com/magwenelab/WeavePop_Cneoformans.

The Supplementary Files and Tables are included in the GitHub repositories mentioned and listed in the Supplementary Material.

The sequencing data and reference genomes used were obtained from NCBI and FungiDB, and their accession numbers are listed in the Supplementary Material.

The authors affirm that all data necessary for confirming the conclusions of the article are present within the article, figures, tables, and the linked materials described above.

## Acknowledgements

The research reported in this publication was supported by the National Institute of Allergy and Infectious Diseases of the National Institutes of Health under award number R01AI133654. The content is solely the responsibility of the authors and does not necessarily represent the official views of the National Institutes of Health. The funders had no role in study design, data collection and analysis, decision to publish, or preparation of the manuscript.

## Supplementary Material

### Analysis of *Cryptococcus neoformans* datasets

#### Analysis of the dataset

The published datasets of short sequencing reads from Desjardins et al. (2017) (387 samples from BioProject PRJNA382844) and Ashton et al. (2019) (884 samples from BioProjects PRJEB27222 and PRJEB5282) (see Table S1) were obtained from NCBI’s Sequence Read Archive. EntrezDirect (Kans 2013, version 16.2) was used to get the run and sample IDs, and the SRA Toolkit (https://github.com/ncbi/sra-tools, version 3.0.10) was used to download the FASTQ files. One sample from the Ashton dataset was not available for download, and 21 samples did not belong to the *C. neoformans* VNI lineage and were excluded from all analyses. The samples with multiple runs were concatenated into a single file. The reads were cleaned with FastP (Chen et al. 2018, version 0.24.0), with default parameters and the `dedup` option.

The reference genomes used were FungiDB release 65 (strain H99, lineage VNI), GCA_022832995.1 (strain VNII, lineage VNII), PRJNA1081442 (strain Bt22, lineage VNBI), and GCA_023650575.1 (strain Bt89, lineage VNBII). To ensure consistent naming, the reference genomes of lineages VNII, VNBI, and VNBII were aligned to the H99 genome using MashMap (Jain et al. 2018, version 3.13) to establish approximate chromosomal homology between them. See Table S3 for correspondence between the reference accessions. The VNI reference annotation (FungiDB release 65) was used to annotate the rest of the reference genomes in the workflow.

We ran WeavePop with parameters listed in the configuration files (Files S8 and S9). From the Ashton dataset, 9 samples had low mapping quality and were removed from further analysis. Additionally, 30 samples (8 Desjardins, 22 Ashton) have depth profiles that suggest a ploidy different than haploid and were also removed. The final results of each dataset (379 samples from Desjardins and 646 samples from Ashton) were combined using the join datasets workflow to leave a final dataset of 1,025 samples.

#### Detection of deletions and duplications in *C. neoformans*

To identify fully and partially deleted and duplicated chromosomes, we used the WeavePop output table, `cnv_chromosomes.tsv`, which contains the percentage of each chromosome covered by each type of CNV. Chromosomes that were 80% or more covered by deletions or duplications were classified as fully deleted or duplicated, respectively. Chromosomes whose CNV coverage was between 20% and 80% were classified as partially deleted or duplicated. And those with less than 20% of coverage by CNVs were deemed euploid (Table S4).

#### Phylogeny

To create an informal supertree with all the analyzed strains, we merged the trees published with the original datasets. The phylogeny by Desjardins et al. (2017) presents the topology among the four lineages: VNI, VNII, VNBI, and VNBII. The tree by Ashton et al. (2019) has only the VNI lineage, but includes the strains published by them as well as the VNI strains in the Desjardins dataset. Thanks to this, we could completely substitute the VNI clade in the Desjardins tree with the full Ashton tree. We rooted the Desjardins tree in the middle of the branch leading to the VNII clade, and the Ashton tree in the middle of the branch leading to the clade VNIa. We removed the VNI clade from the Desjardins tree and used it as the backbone. The Ashton tree was attached to it as a sister clade to the VNB (VNBI and VNBII) lineage. This was done using the function tree.merger (Castiglione et al. 2022) of the R library RRphylo.

### Supplementary Figures

**Figure S1.**
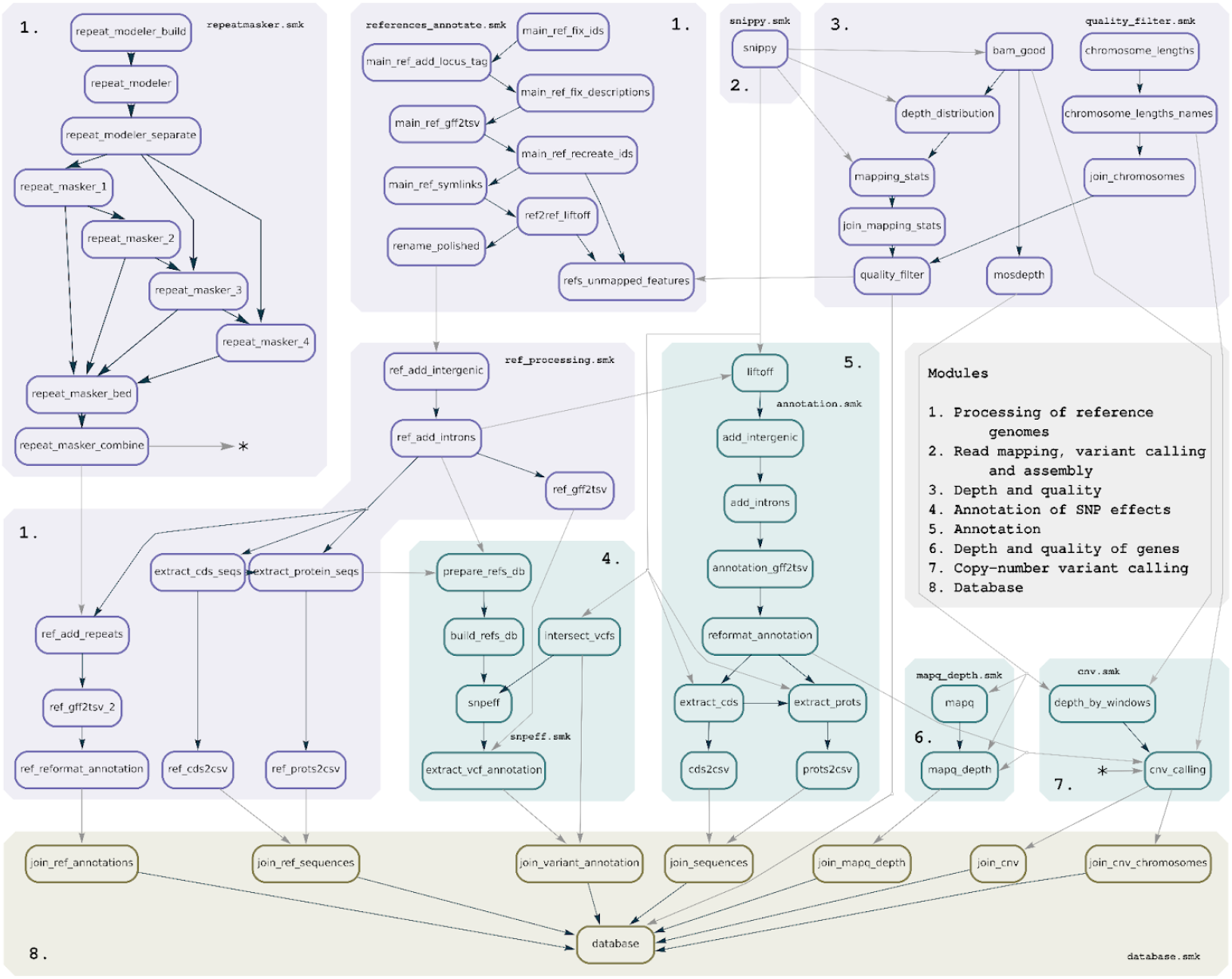
Directed Acyclic Graph (DAG) of jobs in the WeavePop workflow with the Database module and the annotation of reference genomes activated. The trivial rules `ref_fasta_symlinks` and `agat_config` were omitted for simplicity. The rules are located in boxes corresponding to the files where they are defined. The required modules are shown in purple, the intermediate optional ones are in teal, and the database module is in brown.

**Figure S2.**
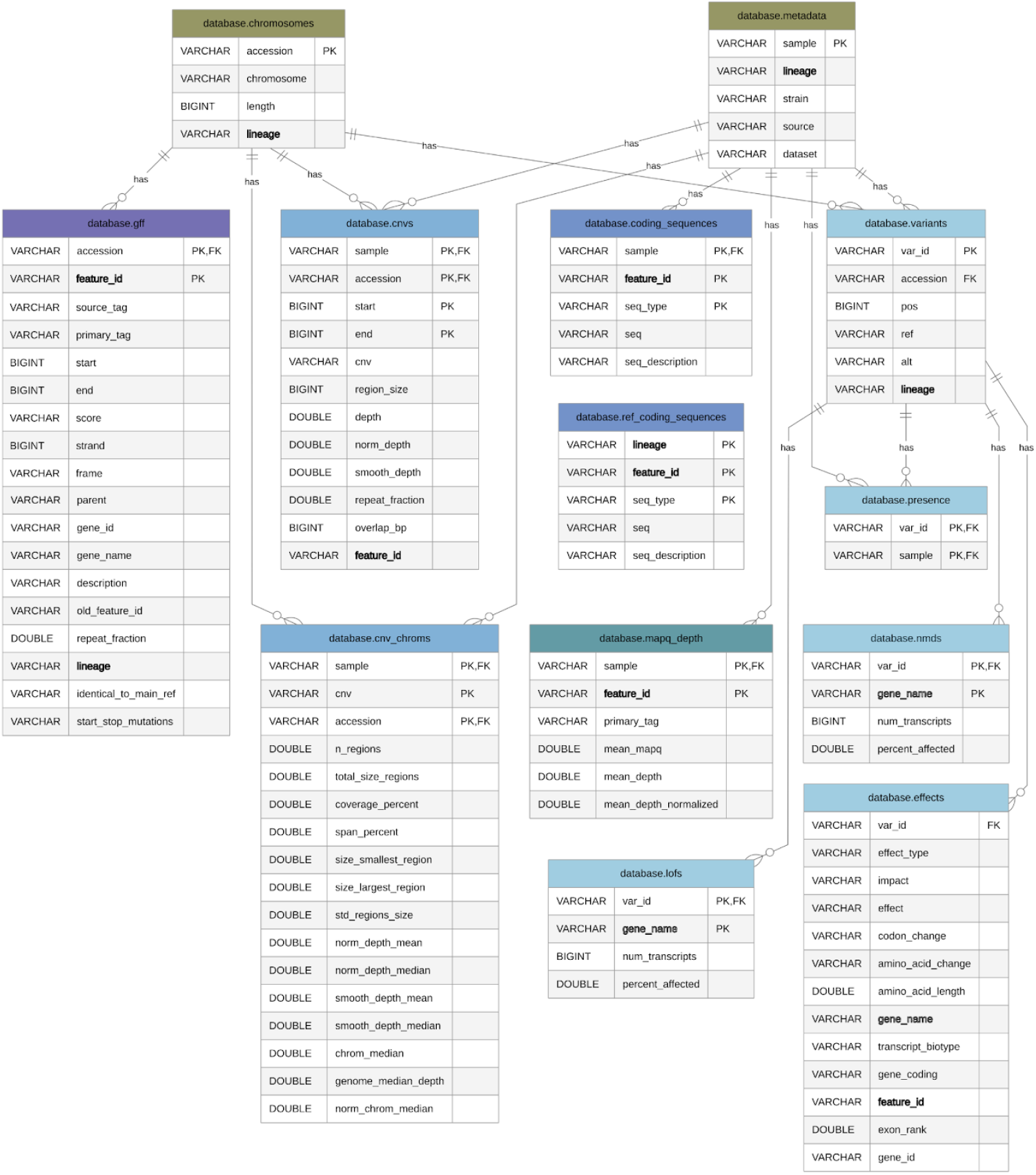
Entity relationship diagram of the database generated by WeavePop. The columns that identify each observation on their own or in combination are defined as primary keys (PK), and the columns that refer to a primary key of another table are defined as foreign keys (FK). Other columns that are common to many tables but are not a unique primary key in any of them are in bold. The tables that are created by the same module have the same color.

**Figure S3.**
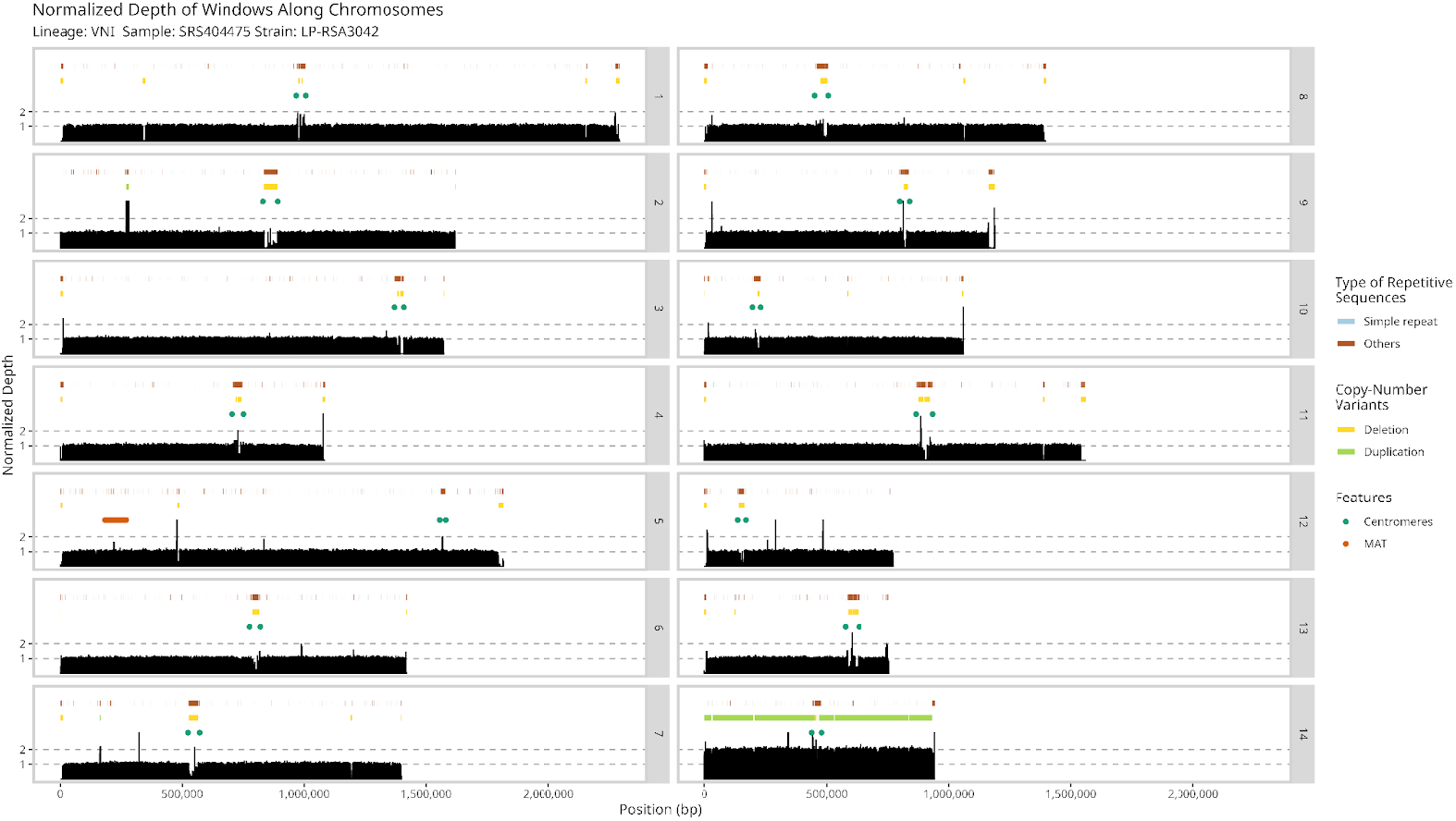
Plot generated by WeavePop showing the normalized read depth of 500 bp windows along all chromosomes of the strain LP-RSA3042. Above the bars showing the read depth, the top track shows segments covered by repetitive sequences in the corresponding reference, the second one shows the identified CNV regions, and the third one shows the location of the genes flanking the centromeres and the mating-type locus. Chromosome 14 is fully duplicated.

**Figure S4.**
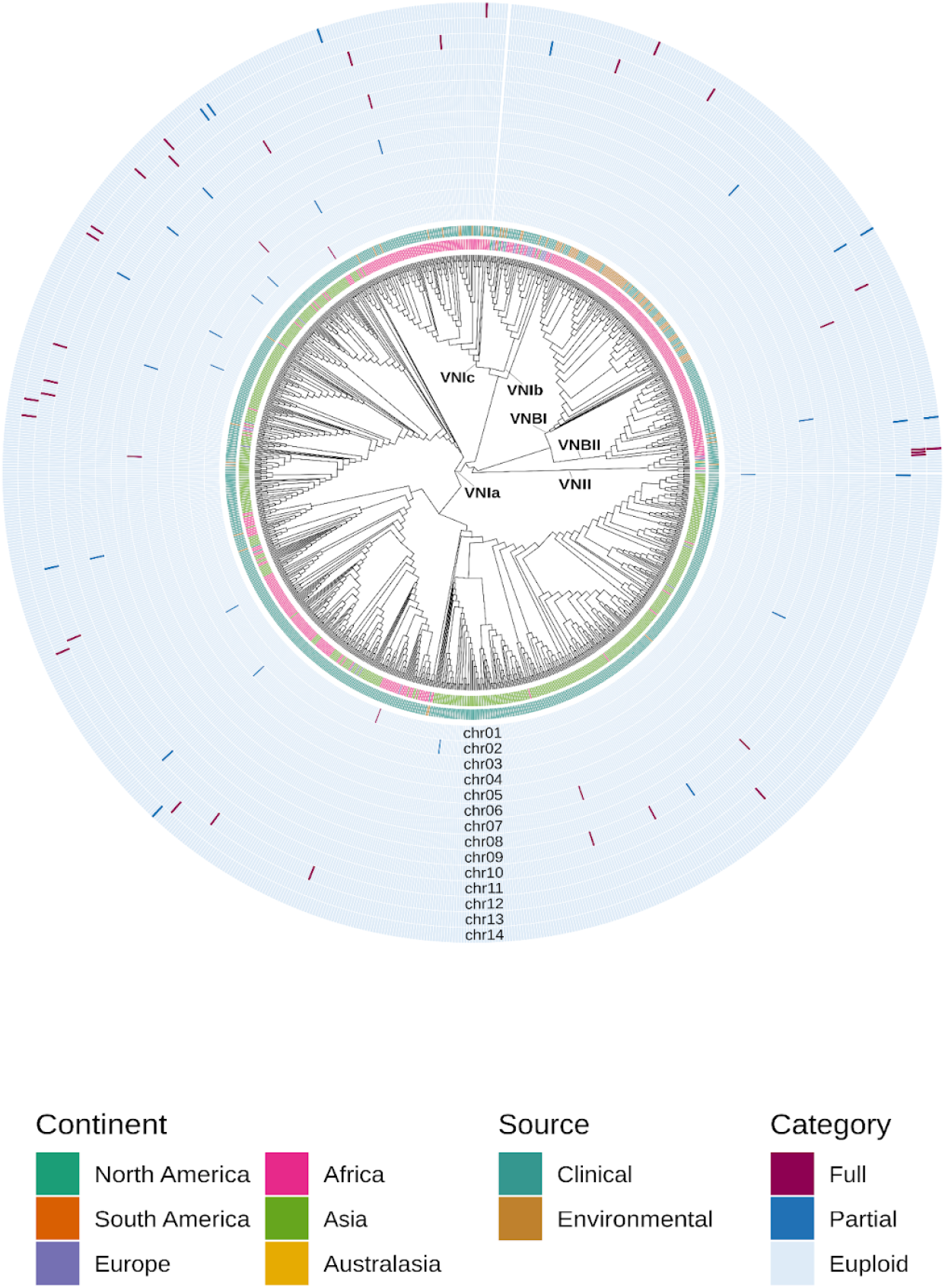
Phylogeny of *C. neoformans* with fully and partially duplicated chromosomes in 1,025 analyzed strains and the reference strain H99. The outer section has one ring per chromosome (in ascending order from the center) with cells colored by category of duplication (Full: >=80% of chromosome length covered by called duplications; Partial: 20-80%; Euploid: 0-20%). The cladogram was constructed by replacing the VNI clade in the phylogeny of Desjardins et al. (2017) with the phylogeny of Ashton et al. (2019) (which includes the VNI Desjardins samples).

### Supplementary Files and Tables

File S1: Input metadata table example: https://github.com/magwenelab/WeavePop/blob/main/test/config/metadata.csv

File S2: Input chromosomes table example: https://github.com/magwenelab/WeavePop/blob/main/test/config/chromosomes.csv

File S3: Input loci table example: https://github.com/magwenelab/WeavePop/blob/main/test/config/loci.csv

File S4: Configuration file: https://github.com/magwenelab/WeavePop/blob/main/config/config.yaml

File S5: Execution profile (YAML with Snakemake command-line options): https://github.com/magwenelab/WeavePop/blob/main/config/default/config.yaml

File S6: WeavePop-CLI examples: https://github.com/magwenelab/WeavePop/wiki/WeavePop%E2%80%90CLI

File S7: Wiki with description of the output: https://github.com/magwenelab/WeavePop/wiki/Output

File S8: Configuration file used to analyze the Ashton dataset: https://github.com/magwenelab/WeavePop_Cneoformans/blob/main/Crypto_Ashton/config/config.yaml

File S9: Configuration file used to analyze the Desjardins dataset: https://github.com/magwenelab/WeavePop_Cneoformans/blob/main/Crypto_Desjardins/config/config.yaml

Table S1: Metadata table of all analyzed samples: https://github.com/magwenelab/WeavePop_Cneoformans/blob/main/analyses/data/processed/metadata_ashton_desj_all_weavepop_final_H99.csv

Table S2: Summary metrics per lineage: https://github.com/magwenelab/WeavePop_Cneoformans/blob/main/analyses/results/tables/per_lineage_summary_stats.tsv

Table S3: Table with correspondence between accessions and chromosome names: https://github.com/magwenelab/WeavePop_Cneoformans/blob/main/Crypto_Desjardins/config/chromosomes.csv

Table S4: Table with the CNV metrics per chromosome of all analyzed samples and their aneuploidy category: https://github.com/magwenelab/WeavePop_Cneoformans/blob/main/analyses/results/tables/chromosome_cnv_categories.tsv

